# ADAM17 Deletion Protects Against Type1 Diabetes-Associated Kidney Injury by Modulating Inflammatory and Fibrotic Pathways

**DOI:** 10.64898/2026.02.05.704041

**Authors:** Marta Riera, Claudia Martyn, Jordi Pujol-Brugués, Eva Marquez, Eva Rodríguez, Vanesa Palau, María José Soler, Juan Sebastián Salazar Castañeda, Melissa Pilco, Jimena del Risco, Marta Crespo, Clara Barrios

**Affiliations:** Hospital del Mar Research Institute, Barcelona, Spain; Nephrology Department Hospital Universitari Vall D’Hebron, Barcelona, Spain; Nephrology Department Hospital del Mar, Barcelona, Spain

## Abstract

0

**Background:** A Disintegrin and Metalloprotease 17 (ADAM17) is a key sheddase regulating multiple inflammatory and growth factor–related pathways implicated in diabetic kidney disease (DKD). While cell-specific deletion of ADAM17 has shown renoprotective effects, the impact of global ADAM17 ablation in the context of diabetes remains incompletely understood.

**Methods:** We investigated the effects of tamoxifen-induced global Adam17 deletion in a murine model of type 1 diabetes induced by streptozotocin. Renal function, structural injury, inflammatory responses, stress-related signalling pathways, and fibrotic remodelling were comprehensively assessed and compared between diabetic Adam17 knockout and control mice.

**Results:** Despite persistent hyperglycaemia and albuminuria, diabetic Adam17 knockout mice exhibited preservation of glomerular filtration rate and marked attenuation of diabetes-associated renal injury. Global Adam17 deletion reduced mesangial expansion and structural damage, limited macrophage infiltration and chemokine expression, and significantly attenuated fibrotic remodelling. At the molecular level, Adam17 deficiency was associated with selective modulation of stress-related signalling pathways, including reduced activation of the PI3K/Akt axis and partial preservation of mitochondrial stress regulators, without evidence of a generalized suppression of cellular stress responses.

**Conclusions:** Our findings demonstrate that global deletion of ADAM17 confers robust protection against diabetes-induced kidney injury through coordinated attenuation of inflammatory activation, stress-related signalling, and fibrotic progression. These results highlight the context-dependent role of ADAM17 in diabetic kidney disease and support the concept that therapeutic strategies targeting ADAM17-related pathways may require tissue- and disease-specific modulation to achieve renoprotective effects.

## 1. Introduction

Diabetic kidney disease (DKD) remains one of the leading causes of chronic kidney disease and end-stage renal disease worldwide, despite significant advances in glycaemic control and the introduction of novel renoprotective therapies [1–4]. Although albuminuria has traditionally been considered an early marker of DKD, progressive loss of glomerular filtration rate (GFR), structural damage, and tubulointerstitial fibrosis ultimately determine renal prognosis [1,5]. Importantly, dissociation between albuminuria and decline in renal function is increasingly recognised, highlighting the complexity of the mechanisms driving disease progression and the need to better understand the molecular pathways underlying diabetic renal injury [1,5].

Chronic inflammation, maladaptive cellular stress responses, and progressive fibrotic remodelling are central features of DKD pathophysiology [4,6]. In both experimental models and human disease, sustained activation of inflammatory signalling pathways promotes chemokine production, recruitment of inflammatory cells, and amplification of profibrotic responses within the kidney [3,4,6]. In parallel, alterations in intracellular survival signalling, mitochondrial dysfunction, and oxidative stress contribute to tubular injury, podocyte loss, and extracellular matrix accumulation, ultimately leading to irreversible renal fibrosis [4,7–9].

A disintegrin and metalloproteinase 17 (ADAM17), also known as tumour necrosis factor-α converting enzyme (TACE), is a membrane-bound sheddase that regulates the proteolytic release of multiple cytokines, growth factors, and receptors involved in inflammatory and growth-related signalling [10,11]. Through cleavage of substrates such as TNF-α, epidermal growth factor receptor (EGFR) ligands, and cytokine receptors, ADAM17 acts as a central regulator of inflammatory and profibrotic pathways [12–16]. Increased ADAM17 expression and activity have been reported in human diabetic nephropathy and experimental models of kidney disease, supporting its relevance in renal pathology and its potential as a therapeutic target [13,17,18].

Our group has previously demonstrated that cell-specific deletion of ADAM17 in renal intrinsic cells confers protection against diabetic kidney injury. In particular, selective ablation of ADAM17 in proximal tubular epithelial cells or endothelial cells attenuated renal inflammation, fibrosis, and functional decline in experimental models of diabetes and metabolic disease [19– 21]. These studies established ADAM17 as a critical mediator of kidney injury within specific renal compartments. However, DKD is a multifactorial disease involving complex interactions between resident renal cells, inflammatory processes, and sustained metabolic stress, and the global contribution of ADAM17 to these integrated mechanisms remains incompletely understood.

In addition to its role in intrinsic renal cells, ADAM17 has been implicated in the regulation of inflammatory responses that shape kidney injury progression. In DKD, enhanced chemokine signalling and macrophage accumulation within the renal cortex contribute to sustained inflammation and fibrotic remodelling [6,22–24]. Whether global inhibition of ADAM17 can attenuate macrophage recruitment and associated inflammatory pathways in the diabetic kidney, thereby limiting downstream structural damage, remains incompletely defined.

In this study, we investigated the impact of complete ADAM17 deletion in a tamoxifen-inducible global knockout mouse model of streptozotocin-induced type 1 diabetes. By integrating functional, histological, and molecular analyses, we aimed to determine whether global ADAM17 deficiency protects against diabetes-associated renal injury and to define the contribution of ADAM17 to inflammatory activation, intracellular stress signalling, and fibrotic remodelling in diabetic nephropathy.

## 2. Materials and Methods

### Animals and generation of global Adam17 knockout mice

A tamoxifen-inducible global Adam17 knockout (Adam17_KO) mouse model was generated by crossing Adam17flox/flox mice (kindly provided by Dr. Raines, Washington University, Saint Louis, MO, USA) with a murine line carrying a Cre recombinase-Estrogen Receptor fusion gene under the control of β-Actin promoter (β-Actin–Cre-ER) obtained from the Jackson Laboratory (Bar Harbor, ME, USA), allowing inducible deletion of Adam17 in all cell types. Adam17flox/flox mice carried loxP sites flanking exon 5 of the Adam17 gene [25], and Cre-mediated recombination resulted in excision of this exon, producing a non-functional transcript. All mice were genotyped using classical PCR technique in DNA isolated from ear clippings. Recombination was induced by tamoxifen administration in 10-week-old mice. Wild-type (WT) littermates lacking loxP sites but receiving identical tamoxifen treatment were used as controls. Successful Adam17 deletion was confirmed in renal tissue by qPCR. All experiments were performed in male mice on a C57BL/6 background. Animals were housed under standard conditions with free access to food and water. All experimental procedures were conducted in accordance with institutional and national guidelines for animal care and use (CEA-OH/10652).

### Experimental design and induction of diabetes

Type 1 diabetes was induced in 12-week-old mice by two intraperitoneal injections of streptozotocin (STZ; 150 mg/kg body weight) administered one week apart to 4 h-fasted animals [26]. Diabetes induction was confirmed by non-fasting blood glucose measurements, and mice with blood glucose levels exceeding 250 mg/dL during the first four weeks after STZ administration were considered diabetic. Diabetic and non-diabetic mice were followed for 20 weeks after diabetes induction. After this period, animals were euthanized under deep anaesthesia [26], blood samples were collected by cardiac puncture, and kidneys were perfused with cold phosphate-buffered saline prior to removal. Renal tissue was processed for histological and molecular analyses.

### Assessment of renal function and metabolic parameters

Renal function was evaluated at the end of the study by measuring glomerular filtration rate (GFR) using fluorescein isothiocyanate (FITC)-inulin clearance following a single bolus injection. GFR values were expressed as μL/min/g body weight [27]. Albuminuria was assessed in morning spot urine samples collected during the final week of follow-up. Urinary albumin concentrations were measured by ELISA (Albuwell M, Exocell, Philadelphia, PA, USA) and creatinine levels were determined using a colorimetric assay (Creatinine Companion, Exocell, Philadelphia, PA, USA). The albumin-to-creatinine ratio (ACR) was calculated and expressed as μg albumin per mg creatinine [28]. Investigators were blinded to genotype during functional assessments.

### Histological and immunohistochemical analyses

Kidney tissues were fixed in formalin and embedded in paraffin for histological evaluation. Periodic acid–Schiff (PAS) staining was performed to assess glomerular morphology and mesangial matrix expansion. The mesangial index was calculated as the ratio of mesangial area to total glomerular tuft area using ImageJ software [29]. Immunohistochemical analyses were performed to evaluate podocyte number (WT1), macrophage infiltration (F4/80), myofibroblast activation (α-smooth muscle actin, α-SMA), and fibrotic remodelling (galectin-3). Positive staining was quantified in a predefined number of fields per animal, and analyses were conducted in a blinded manner.

### Gene expression analysis

Total RNA was isolated from renal cortex tissue with Tripure Isolation Reagent (Sigma-Aldrich, Saint Louis, MO, USA) and reverse-transcribed into cDNA using the High Capacity cDNA RT Kit (ThermoFisher Scientific, Waltham, MA, USA) [30]. Gene expression levels of Adam17, monocyte chemoattractant protein-1 (Mcp1/Ccl2), and chemokine (C–C motif) ligand 5 (Ccl5) were quantified by real-time PCR using LightCycler®480 SYBR Green I Master Mix (Roche, Basel, Switzerland). Hypoxanthine phosphoribosyltransferase 1 (Hprt) was used as a housekeeping gene for normalization. Relative gene expression was calculated using the 2^−ΔΔCt^ method.

### Protein expression analysis

Protein extracts from renal cortex tissue homogenized in extraction buffer containing 50 mM HEPES, pH 7.4, 150 mM NaCl, 0.5% Triton X-100, 0.025 mM ZnCl2, (all from Sigma-Aldrich, Saint Louis, MO, USA) 0.1 mM Pefabloc SC Plus (Roche, Basel, Switzerland), EDTA-free protease inhibitor cocktail tablet (Roche, Basel, Switzerland), and phosphatase inhibitor cocktail (Sigma-Aldrich, Saint Louis, MO, USA) were analyzed by Western blotting. Fifteen μg of total protein were used to assess expression of phosphorylated and total Akt, MCP-1, galectin-3, SIRT3, and FoxO3. Protein expressions were quantified by pixel intensity and normalized to appropriate loading controls.

### Tumor necrosis factor-α measurements

Tumor necrosis factor-α (TNF-α) levels were measured in serum and renal cortex homogenates using a commercially available ELISA kit according to the manufacturer’s instructions Kit (R&D Systems, Minneapolis, MN, USA). TNF-α was assessed as an exploratory inflammatory marker.

### Statistical analysis

Data are presented as mean ± standard error (SE). Statistical analyses were performed using one-way ANOVA for normally distributed data or non-parametric Kruskal–Wallis and Mann– Whitney tests when appropriate. A p value < 0.05 was considered statistically significant.

## 3. Results

### 3.1 Validation of tamoxifen-induced global Adam17 deletion

Tamoxifen administration efficiently induced global deletion of Adam17 in adult mice. Adam17 gene expression was markedly reduced in renal tissue from Adam17 knockout (KO) mice compared with wild-type (WT) controls, confirming effective recombination (Figure 1B). Baseline metabolic parameters and body weight did not differ between genotypes prior to diabetes induction, indicating that global Adam17 deletion did not affect general metabolic status under non-diabetic conditions. At study completion, both diabetic groups, WT and KO, displayed comparable hyperglycemia, significantly elevated relative to non-diabetic control (Figure 1C).

**Figure 1.**
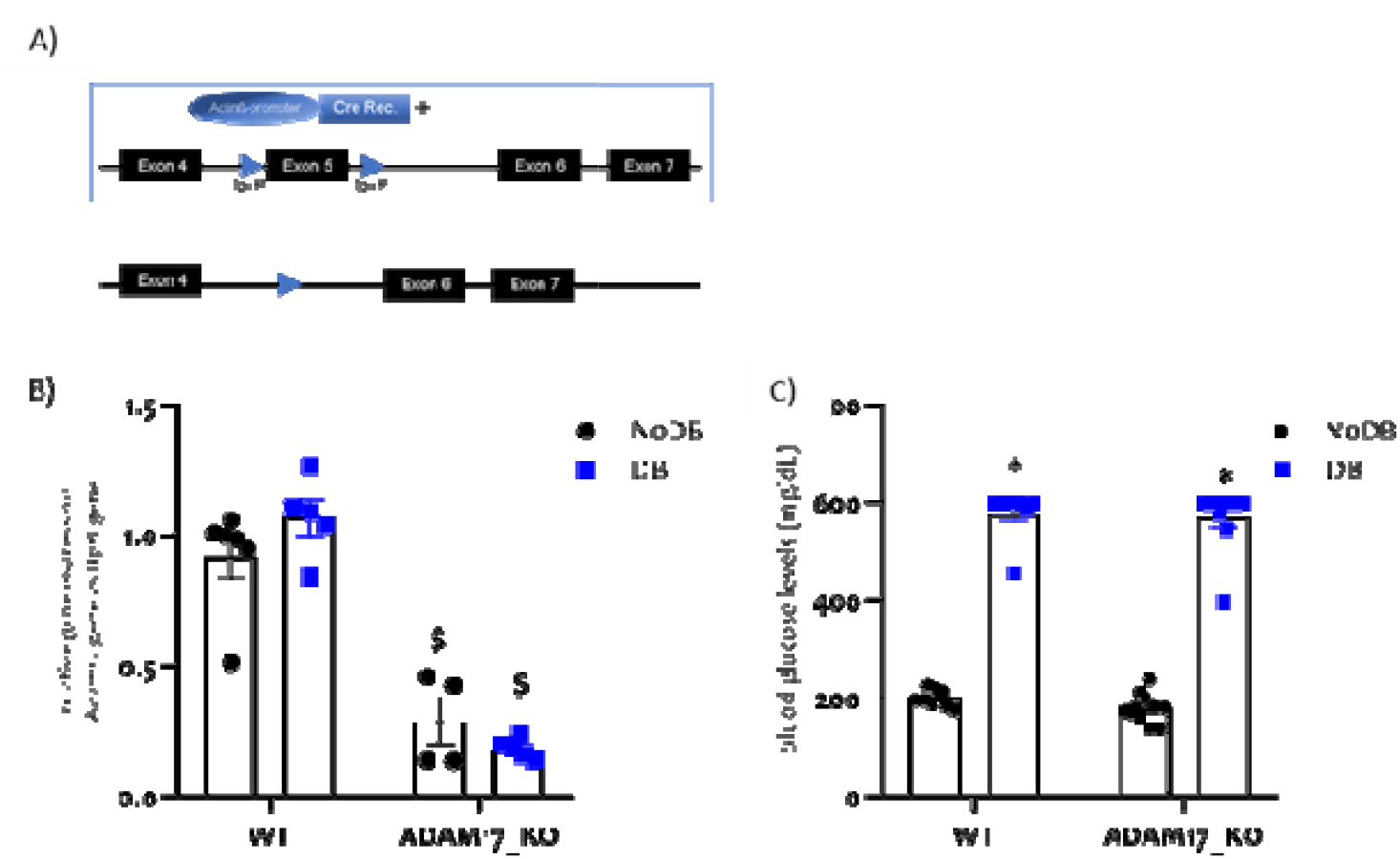
Validation of tamoxifen-induced global Adam17 deletion. (A) Schematic representation of the tamoxifen-inducible global Adam17 knockout model. (B) Adam17 mRNA expression levels in renal tissue, confirming effective recombination in Adam17 knockout (KO) mice compared with wild-type (WT) controls. (D) Final blood glucose levels. STZ-treated mice remained diabetic throughout the 20-week follow-up. Data are presented as mean ± SEM. Groups: non-diabetic (NoDB); diabetic (DB); wild-type (WT); Adam17 knockout (ADAM17_KO). *p≤0.05 vs. NoDB; $p≤0.05 vs. WT.

### 3.2 Global Adam17 deletion preserves renal function despite persistent albuminuria

To evaluate the impact of global Adam17 deletion on renal function in diabetes, glomerular filtration rate (GFR) and albuminuria were assessed after 20 weeks of streptozotocin-induced diabetes. Diabetic WT mice exhibited a significant decline in GFR compared with non-diabetic controls. In contrast, diabetic Adam17 KO mice preserved GFR, showing significantly higher filtration rates than diabetic WT animals (Figure 2A).

**Figure 2.**
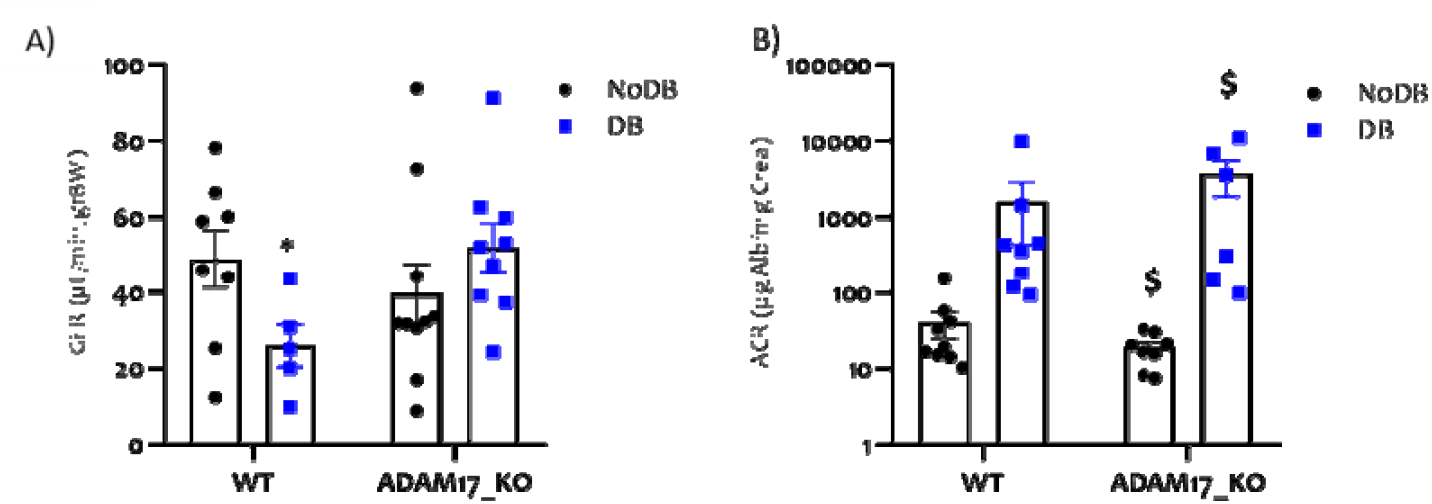
Global Adam17 deletion preserves renal function despite persistent albuminuria. (A) Glomerular filtration rate (GFR) measured after 20 weeks of diabetes. (B) Albumin-to-creatinine ratio (ACR) in a logarithmic scale, showing increased albuminuria independently of genotype. Data are presented as mean ± SEM; 5-10 animals were included per group. Groups: non-diabetic (NoDB); diabetic (DB); wild-type (WT); Adam17 knockout (ADAM17_KO). *p≤0.05 vs. NoDB; $p≤0.05 vs. WT.

Albuminuria was markedly increased in both diabetic WT and Adam17 KO mice, with no significant differences between genotypes (Figure 2B). These findings indicate a dissociation between albuminuria and renal functional decline, suggesting that global Adam17 deletion preferentially modulates mechanisms related to loss of filtration capacity rather than glomerular permeability.

### 3.3 Adam17 deletion attenuates diabetes-induced structural kidney damage

Consistent with preserved renal function, histological analyses revealed that global Adam17 deletion attenuated diabetes-induced structural kidney injury. Periodic acid–Schiff (PAS) staining demonstrated significant mesangial expansion in diabetic WT mice, which was markedly reduced in diabetic Adam17 KO animals (Figure 3A and 3C). Furthermore, podocyte loss, assessed by WT1 immunostaining, was evident in diabetic WT kidneys but was significantly attenuated in Adam17-deficient mice (Figure 3B and 3C), indicating preservation of glomerular architecture.

**Figure 3.**
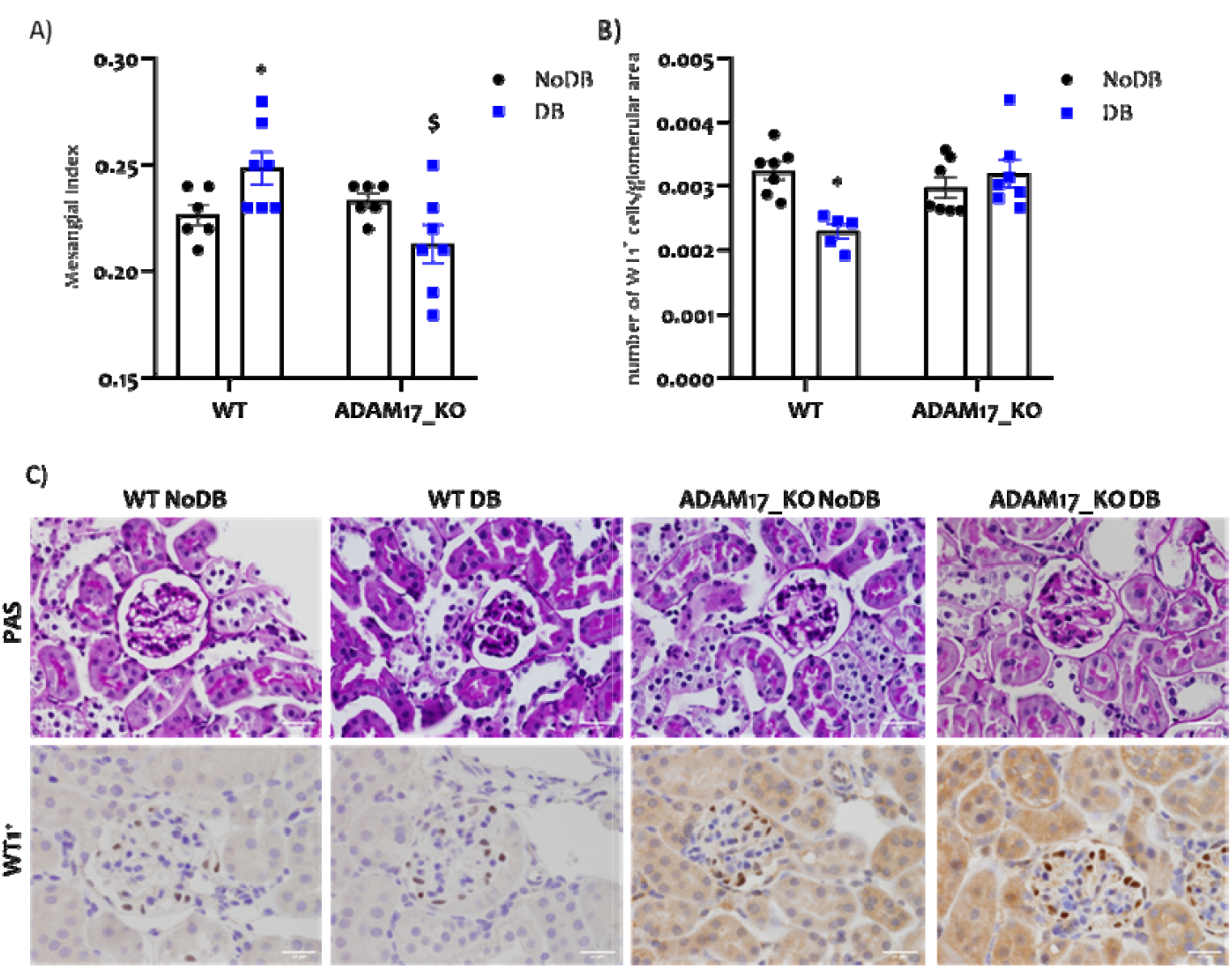
Adam17 deletion attenuates diabetes-induced structural kidney damage. (A) Mesangial expansion quantification from PAS staining samples. (B) Podocytes detected by WT1 staining, quantified, and normalized to glomerular area. (C) Representative images for PAS- and WT1-stained images at 400x magnification (scale bar = 50μm). Data are presented as mean ± SEM. *p≤0.05 vs. NoDB; $p≤0.05 vs. WT.

### 3.4 Reduced macrophage infiltration and chemokine-associated inflammation in Adam17 KO kidneys

Renal inflammatory activation was evaluated by assessing macrophage infiltration and chemokine expression. Diabetic WT mice exhibited pronounced accumulation of F4/80-positive macrophages within the renal cortex. In contrast, diabetic Adam17 KO mice showed similar levels as non-diabetic Adam17 KO mice (Figure 4A–B). At the molecular level, type 1 diabetes induced an increase in *Ccl5* gene expression in WT mice (Figure 4C). Surprisingly, non-diabetic Adam17-KO mice displayed similarly elevated basal Ccl5 levels compared with diabetic Adam17 KO mice, suggesting a diabetes-independent inflammatory tone associated with Adam17 deficiency. Expression of *Mcp1* (*Ccl2*) gene was significantly increased in diabetic WT kidneys, while the differences between diabetic and non-diabetic KO mice were not statically significant. (Figure 4D). These results were better delineated at protein level where all Adam17 KO mice showed similar expression levels despite the diabetic state (Figure 4D-E). Together, these data indicate that global Adam17 deletion limits macrophage-associated inflammatory activation related to Mcp1.

**Figure 4.**
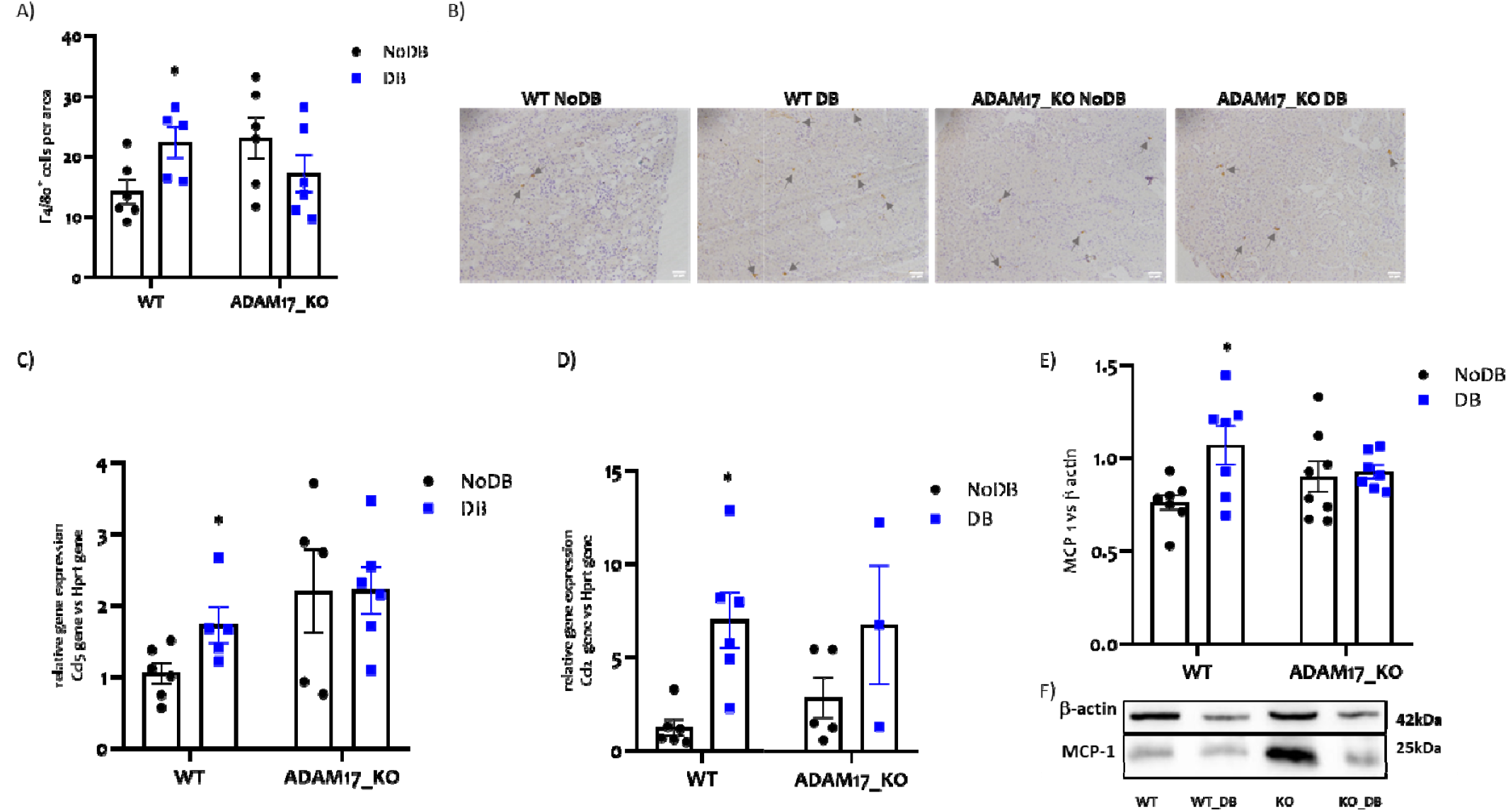
Reduced macrophage infiltration and chemokine-associated inflammation. (A) F4/80^+^-macrophages quantified and normalized to glomerular area; (B) representative images of cell infiltration at 200x magnification where grey arrows indicate F4/80^+^ cells; (C) Ccl5 expression after 20 weeks of follow up; (D) renal Ccl2/MCP-1 gene expression; (E) MCP-1 protein levels normalized to β-actin; (F) representative Western blot images. Data are presented as mean ± SEM. Groups: non-diabetic (NoDB); diabetic (DB); wild-type (WT); Adam17 knockout (ADAM17_KO). *p≤0.05 vs. NoDB.

Furthermore, given the role of ADAM17 in TNF-α shedding, we wanted to asses TNF-α levels as an exploratory inflammatory marker. TNF-α concentrations in serum as the shedded form and the ones in renal tissue showed no consistent results between diabetic WT and Adam17 KO mice (Figure S1).

### 3.5 Selective modulation of stress-related signalling pathways in Adam17-deficient kidneys

To explore intracellular signalling mechanisms associated with Adam17 deletion, key pathways involved in cellular stress and survival were analysed. Diabetes induced marked activation of the PI3K/Akt signalling pathway in WT kidneys, as evidenced by increased Akt phosphorylation. This response was significantly attenuated in diabetic Adam17 KO mice where both groups had similar values (Figure 5A–B).

**Figure 5.**
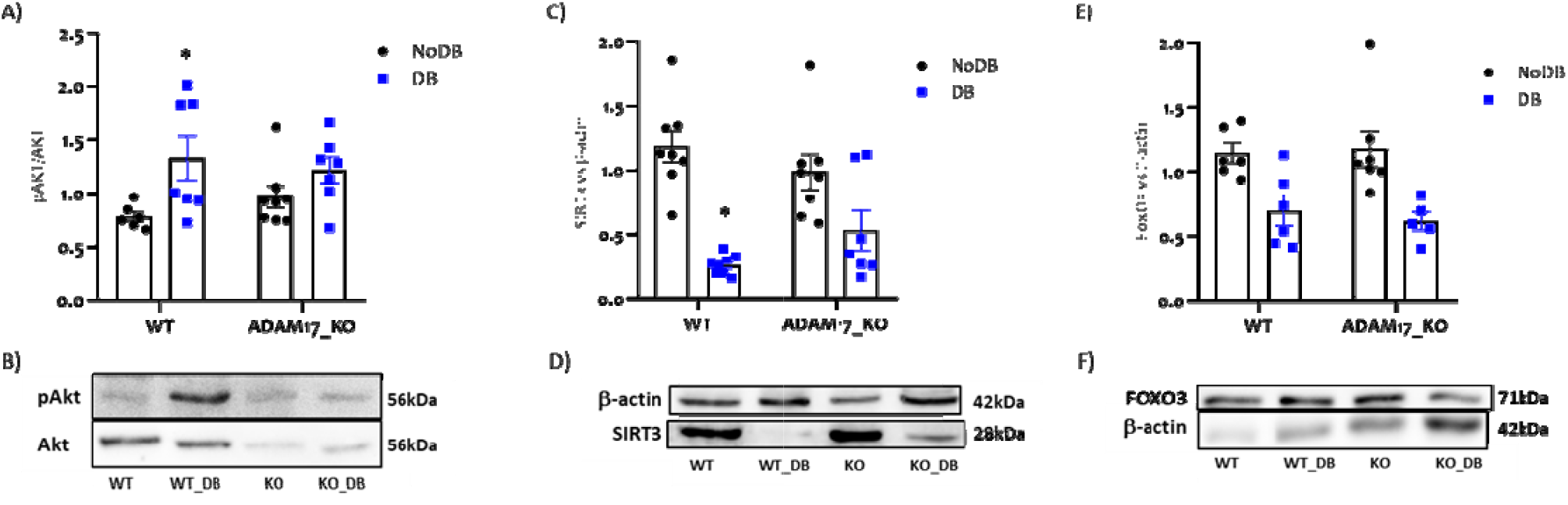
Selective modulation of stress-related signaling pathways. (A–B) p-Akt/Akt analysis and representative Western blot images for pAkt and Akt; (C–D) SIRT3 protein normalized to β-actin and representative Western blot images; (E-F) FOXO3 protein bands normalized to β-actin and representative Western blot images. Data are presented as mean ± SEM. Groups: non-diabetic (NoDB); diabetic (DB); wild-type (WT); Adam17 knockout (ADAM17_KO). *p≤0.05 DB vs NoDB.

Analysis of mitochondrial and oxidative stress regulators revealed a more selective pattern of modulation. SIRT3 expression was partially preserved in Adam17-deficient kidneys compared with diabetic WT controls (Figure 5C–D), whereas FoxO3 expression and activation remained largely unchanged (Figure 5E–F).

### 3.6 Global Adam17 deletion limits fibrotic remodeling in diabetic kidneys

Fibrotic remodeling was markedly reduced in diabetic Adam17 KO mice. Immunohistochemical analysis demonstrated significant upregulation of α-smooth muscle actin (α-SMA) and galectin-3 in diabetic WT kidneys, reflecting myofibroblast activation and fibrotic progression. In contrast, expression of both markers was significantly attenuated in Adam17-deficient mice (Figure 6).

**Figure 6.**
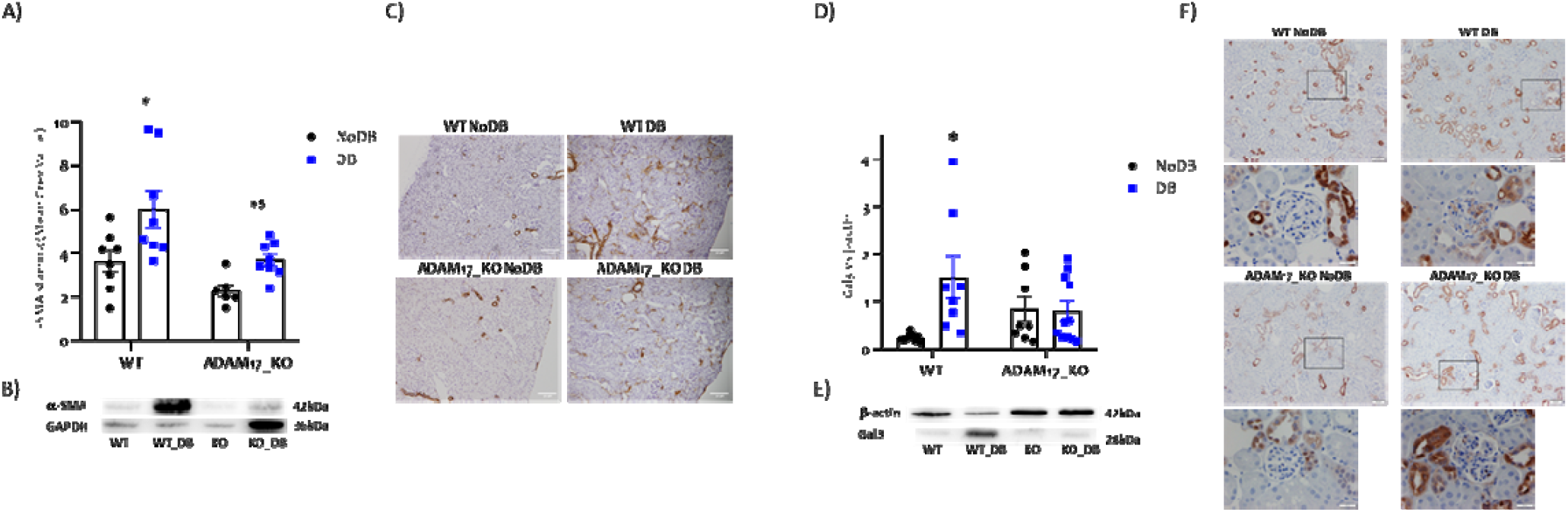
Global Adam17 deletion limits fibrotic remodeling. (A, C) Fibrosis due to α-SMA staining and quantification on cortex kidney sections as shown in the microphotographs at 200x magnification; (B) α-SMA protein bands compared with GAPDH bands in a representative Western blot image; (D-E) Galectin-3 protein bands normalized to β-actin and representative bands in Western blot images; (F) Microphotographies of Gal3 localized by immunohistochemistry at 100x and 400x magnification. Data are presented as mean ± SEM. Groups: non-diabetic (NoDB); diabetic (DB); wild-type (WT); Adam17 knockout (ADAM17_KO). *p≤0.05 vs. NoDB.

Collectively, these results demonstrate that global Adam17 deletion confers significant protection against diabetes-associated kidney injury by preserving renal function, attenuating structural damage, limiting macrophage-associated inflammation, selectively modulating stress-related signalling pathways, and reducing fibrotic remodelling.

## 4. Discussion

In this study, we demonstrate that tamoxifen-induced global deletion of ADAM17 confers significant protection against diabetes-associated kidney injury in a murine model of type 1 diabetes. Despite persistent hyperglycemia and albuminuria, Adam17-deficient mice exhibited preservation of renal function, reduced structural damage, attenuated inflammatory activation, and markedly decreased fibrotic remodeling. These findings identify ADAM17 as a central modulator of pathological pathways driving diabetic kidney disease progression and extend previous observations obtained using cell-specific Adam17 deletion models reported by our group [19–21].

One of the most relevant observations in this study is the dissociation between preservation of glomerular filtration rate and persistent albuminuria in diabetic Adam17-deficient mice. Although albuminuria has traditionally been considered an early marker of diabetic nephropathy, accumulating evidence indicates that albuminuria and progressive loss of renal function reflect partially independent pathophysiological processes [1,5]. In this context, our data suggest that ADAM17 deletion preferentially modulates mechanisms related to structural injury, nephron integrity, and fibrotic progression rather than alterations in glomerular permeability [4,7].

Chronic inflammation plays a central role in the progression of diabetic kidney disease, contributing to both structural damage and fibrotic remodeling [2–4,6]. In the present study, global Adam17 deletion was associated with reduced renal macrophage infiltration and decreased expression of chemokines involved in monocyte recruitment, including MCP-1 and CCL5. Importantly, our findings support a role for ADAM17 in regulating macrophage-associated inflammatory recruitment rather than a generalized immune response. While we did not assess macrophage activation states or functional phenotypes, the reduction in macrophage accumulation is consistent with a diminished inflammatory microenvironment that may limit subsequent tissue injury and fibrogenesis [31–34].

In addition to its effects on inflammatory recruitment, ADAM17 deletion selectively modulated intracellular stress-related signaling pathways. Diabetes-induced activation of the PI3K/Akt pathway, which has been implicated in maladaptive repair responses and fibrotic progression in chronic kidney disease [7,8,35,36], was attenuated in Adam17-deficient kidneys. In contrast, regulators of mitochondrial and oxidative stress exhibited a more nuanced pattern of modulation, with partial preservation of SIRT3 expression and no significant changes in FoxO3 signaling. These findings suggest that ADAM17 does not act as a global regulator of cellular stress responses, but rather modulates specific signaling pathways in a context-dependent manner [9,37,38].

Fibrotic remodeling emerged as one of the most robustly affected processes following global Adam17 deletion. Reduced expression of α-smooth muscle actin and galectin-3 indicates attenuation of myofibroblast activation and extracellular matrix remodeling in diabetic Adam17-deficient mice. Fibrosis represents a final common pathway in chronic kidney disease progression, integrating inflammatory, metabolic, and cellular stress signals [22,39–42]. Our findings therefore support a role for ADAM17 as a molecular link between chronic inflammatory activation and fibrotic progression in the diabetic kidney, providing a mechanistic basis for the observed preservation of renal function [23,24,43,44].

The pleiotropic nature of ADAM17 raises important considerations regarding its role across different tissues and disease contexts. ADAM17 regulates the shedding of multiple substrates involved in inflammation, growth factor signaling, and cell–cell communication, and its functional consequences are highly dependent on on cellular and organ-specific settings [10– 18]. Accordingly, divergent effects of ADAM17 deletion have been reported across experimental models, underscoring the context-dependent actions of this protease [45]. In line with this complexity, previous strategies aimed at systemic or catalytic site inhibition of ADAM17 have faced important limitations, largely related to limited selectivity, tolerability issues, and interference with homeostatic shedding functions, which may contribute to heterogeneous or context-dependent outcomes across disease models [46–48]. Our findings support the concept that therapeutic benefit from ADAM17 modulation is likely context-dependent and may rely on identifying downstream pathways that are particularly relevant in the diabetic kidney. In this regard, the ADAM17–Klotho axis emerges as an attractive mechanistic link, as increasing evidence indicates that ADAM17 can promote Klotho shedding and that disruption of this pathway may influence oxidative stress–related injury and tissue vulnerability in diabetic kidney disease [49]. Finally, while direct inhibition of ADAM17 by sodium–glucose cotransporter 2 inhibitors has not been demonstrated, recent studies have suggested that some of the pleiotropic renoprotective effects of SGLT2 inhibitors may involve modulation of inflammatory and stress-related pathways converging on ADAM17-related signalling including Klotho-associated mechanisms [50,51]. Although this hypothesis remains speculative, it is conceptually aligned with the renoprotective profile observed in our model.

Some limitations of this study should be considered. First, the use of a global Adam17 knockout model precludes attribution of specific effects to individual renal or immune cell populations. Second, inflammatory analyses were limited to macrophage infiltration and chemokine expression, without functional characterization of immune cell phenotypes. Third, this study was performed in a murine model of type 1 diabetes, and extrapolation to other forms of diabetic kidney disease or human pathology should be made with caution. Despite these limitations, the integration of functional, structural, and molecular analyses provides a comprehensive assessment of the impact of ADAM17 deletion on diabetic kidney injury.

In conclusion, our results demonstrate that global deletion of ADAM17 protects against diabetes-associated renal injury by attenuating inflammation, selectively modulating stress-related signaling pathways, and limiting fibrotic remodeling, ultimately preserving renal function. These findings extend previous cell-specific studies and highlight the context-dependent role of ADAM17 in diabetic kidney disease, supporting the need for refined and tissue aware therapeutic strategies targeting this protease.

## Supporting information

Supplemental tables 1 and 2 and suppFigure

