## Supplemental tables 1 and 2 and suppFigure for "ADAM17 Deletion Protects Against Type1 Diabetes-Associated Kidney Injury by Modulating Inflammatory and Fibrotic Pathways"

**SUPPLEMENTARY INFORMATION**

***Supplementary Table 1. List of primers used for the analysis of the genes.*** The table is divided according the type of primers, located at genomic or at cDNA sequences used for both processes.

| **Target DNA** | **Primer Forward** | **Primer Reverse** |
| --- | --- | --- |
| actinCre | 5’ CCTGGCGATCCCTGAACATGTCC 3’ | 5’ CTCTAGAGCCTCTGCTAACC 3’ |
| Adam17Fl_F | 5’ ATAGGGAGCCAAGTGTGATGG 3’ | 5’ CACATACTTGCCTACAAGCCAG 3’ |
| **Target mRNA** | **Primer Forward** | **Primer Reverse** |
| Adam17 | 5’ GGCAGAATATAACGTAGAGCCACT 3’ | 5’ CTTCAGACTTATACACCAGC 3’ |
| Mcp1/Ccl2 | 5’ AGGTCCCTGTCATGCTTCTG 3’ | 5’ CGTTAACTGCATCTGGCTGA 3’ |
| Ccl5 | 5’ CTGCTGCTTTGCCTACCTCT 3’ | 5’ GTGACAAACACGACTGCAAGAT 3’ |
| Hprt | 5’ TGTTGTTGGATATGCCCTTG 3’ | 5’ AATGACACAAACGTGATTCAAA 3’ |

***Supplementary Table 2. Antibodies used for immunohistochemistry and for Western blot (WB).*** Secondary antibodies for the WB were Peroxidase AffiniPure Donkey Anti-Rabbit IgG (H+L) and Peroxidase AffiniPure Donkey Anti-Mouse IgG (H+L) (Jackson Immunoresearch). BSA: Bovine Serum Albumin; GS: Goat Serum; NFM: Non-Fat Milk diluted in TBS-T 0.1%

| **Antibody target** | **Source** | **Cat. Number** | **Company** | **Working dilution** |
| --- | --- | --- | --- | --- |
| **Immunohistochemistry** | | | | |
| F4/80 | rat | 123101 | Biolegend (#123101) | 1:500 in 3%BSA/3%GS |
| WT1 | rabbit | 12609-1-AP | Proteintech | 1:1000 in 3%BSA/3%GS |
| α-SMA | mouse | A-2547 | Sigma-Aldrich | 1:1000 comprovar |
| Galectin3 | mouse | 126701 | Biolegend | 1:500 comprovar |
| **Western Blot** | | | | |
| pAKT (Ser473) | rabbit | 9271 S | Cell Signaling Technology | 1:1000 in 2.5% BSA |
| Akt | rabbit | 9272 S | Cell Signaling Technology | 1:2000 in 2.5% BSA |
| MCP1/CCL2 | mouse | TA336914 | Origene | 1:2000 in 2.5% NFM |
| SIRT3 | rabbit | CAB7307 | Assay Genie | 1:3000 in 2.5% NFM |
| FoxO3 | rabbit | CAB0102 | Assay Genie | 1:3000 in 2.5% NFM |
| α-SMA | mouse | A-2547 | Sigma-Aldrich | 1:1000 in 2.5% NFM |
| Galectin3 | mouse | 126701 | Biolegend | 1:1000 in 2.5% NFM |
| ß-actin | mouse | A1978 | Sigma-Aldrich | 1:20000 in 2.5% NFM |
| Gapdh | mouse | sc-32233 | Santa Cruz Biotechnology | 1:20000 in 2.5% NFM |

***Supplementary Figure 1. Influence of diabetes and Adam17 deletion on circulating and kidney cortex TNF-α***. A) Serum TNF-α quantified by ELISA; B) TNF-α levels in kidney cortex homogenates. Data are presented as mean ± SEM. Groups: non-diabetic (NoDB); diabetic (DB); wild-type (WT); Adam17 knockout (ADAM17_KO).

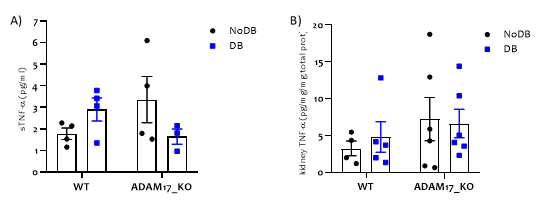
